# Transcriptional analysis shows a robust host response to *Toxoplasma gondii* during early and late chronic infection in both male and female mice

**DOI:** 10.1101/523183

**Authors:** Andrew L. Garfoot, Patrick W. Cervantes, Laura J. Knoll

## Abstract

The long-term host effects caused by the protozoan parasite *Toxoplasma gondii* are poorly understood. RNA-seq analysis previously determined that the host response in the brain was higher and more complex at 28 versus 10 days postinfection. Here, we analyzed the host transcriptional profile of age-and sex-matched mice during early (21 and 28 days) and late (3 and 6 months) chronic infection. We found that a majority of the host genes which increase in abundance at day 21 postinfection are still increased 6 months postinfection for both male and female mice. While most of the differentially expressed genes were similar between sexes, females have far fewer genes that are significantly less abundant, which may lead to the slight increased cyst burden in males. Transcripts for C-X-C Motif Chemokine Ligand 13 (CXCL13) and a C-C Motif Chemokine Receptor 2 (CCR2) were significantly higher in females compared to males during infection. As *T. gondii* chronic infection and profilin (PRF) confer resistance to *Listeria monocytogenes* infection in a CCR2 dependent manner, the sex specific difference in CCR2 expression lead us to re-test the protection of PRF in both sexes. Chronic infection as well as PRF were nearly as effective at reducing the bacterial burden in male versus female mice. These data show that most of the differentially express host genes are similar between males and females, important differences exist leading us to emphasize the inclusion of both sexes for future studies.

## INTRODUCTION

The parasitic protozoan *Toxoplasma gondii* has the unique ability to form long-lasting chronic infections in the central nervous system (CNS) of humans and other mammals. After the initial infection through ingestion of either *T. gondii* oocysts or tissues cysts, the fast-growing tachyzoite form travels into the CNS and transitions into the slow-growing bradyzoite. These bradyzoites form cysts in the brain, primarily within neurons (1), and reside there presumably for the lifetime of the host. The long-term effects of chronic *T. gondii* infection on the body and brain are poorly understood, and current antiparasitic drugs are ineffective against this chronic form. Although otherwise healthy individuals chronically infected with *T. gondii* are clinically asymptomatic, the risk of the risk of cyst reactivation is inherent when an infected person becomes immunocompromised.

In chronically infected mice, *T. gondii* provides protection against lethal pathogens (2–7), largely through a strong immune response against the immunodominant antigen profilin (PRF, (8)). PRF binds the murine Toll-like receptors 11 and 12 (TLR11/12) which leads to an interleukin-12 (IL-12) dependent interferon gamma (IFNγ) response (9–12). Continual IFNγ production is required to suppress chronic *T. gondii* infection, as reduction of IFNγ in chronically infected mice results in tissue cyst reactivation and host death (13, 14). Although the immune mechanism is different in humans, a continuous suppression of *T. gondii* by the immune system is required for survival (15). This effect is evident in patients with chronic *T. gondii* infection who become immune compromised by an immunosuppressive disease or chemotherapeutic treatment. These patients suffer from encephalitis when brain cysts reactivate into tachyzoites.

The immune response to parasitic protozoans, like *T. gondii, Leishmania*, and *Plasmodium*, vary between male and female hosts. During acute *T. gondii* infection, females show reduced survival rates and lower cytokine levels compared to males (16). Innate and adaptive immune cells are influenced by sex hormones that impact the immune response to protozoan infections (17). Treatment with sex hormones, like estradiol and estrogen, increase acute pathogenesis (18) and increase the number of brain cysts in both male and female mice (19, 20). Although the effects of sex hormones on acute *T. gondii* infection have been well studied, comparisons between male and females during chronic infection have varied depending on mouse genotypes. Cyst counts between male and female outbred mice show no difference between sexes (19), whereas inbred B6 female mice have slightly more brain cyst compared to male during early chronic infection (16).

To start understanding the long-term interactions between the parasite and host, the host/parasite transcriptomic profile was compared at the peaks of acute and chronic mouse infection (21). The number of highly expressed host genes specific to chronic infection was striking and required further investigation. In this manuscript, we used high throughput RNA sequencing (RNAseq) to examine the host response late into chronic

*T. gondii* infection. Brain tissue of CBA/J mice infected from 21 days through 6 months of infection was analyzed, and both male and female mice were included to account for sex dependent responses to chronic *T. gondii* infection. We found that a majority of the host genes which are increased in abundance at 21 days postinfection are still more abundant at 6 months postinfection in both sexes. While most of the host response is similar between sexes, females have far fewer genes that are significantly less abundant. Also, the B-cell chemoattractant C-X-C Motif Chemokine Ligand 13 (CXCL13) and a C-C Motif Chemokine Receptor 2 (CCR2) for Monocyte Chemoattractant Protein 1 (MCP1) are higher in females compared to males during infection. These results highlight that studies of the host immune response against *T. gondii* should include both males and females, especially when examining responses to the chemokines and their receptors.

## RESULTS

### Host response to *T. gondii* throughout chronic infection

We sequenced the transcripts from infected cerebral cortex tissue in mice to examine the host expression dynamics during both early (21 and 28 days postinfection) and late (90 and 180 days postinfection) chronic *T. gondii* infection (Fig. 1). Uninfected age-and sex-matched control mice were sequenced at each timepoint (n=2-3) to account for age-and sex-related changes independent of infection (Fig. 1A). To reduce sequencing bias, RNA libraries were multiplexed and run across 6 lanes for infected tissues and 2 lanes for uninfected tissues. Sequencing generated approximately 46 million paired-end, 125 base pair reads per lane for each sample, providing approximately 270 million reads for each infected tissue sample and 100 million reads for uninfected tissue sample. Reads were aligned to the *Mus musculus* genome (normalized expression values are listed in Table S1) for analysis. Reads were also aligned to *T. gondii* (normalized values in Table S2); however, the RNA processing methods used did not allow for efficient RNA extraction from the cysts, so no comparisons were performed with the *T. gondii* transcripts. Analyzing the similarity of samples using the mouse alignment data through a principle component analysis showed distinct grouping between male and female groups as well as infected and uninfected samples (Fig. 1B). Within each group, separation of the 180 day samples is seen, likely due to age related changes in the mice since this happens in both uninfected and infected samples. Pearson’s correlation coefficients show high similarity among all samples (ranging from 84-99% similar; Table S3), with the greatest difference between infected and uninfected samples of either sex.

**Fig. 1.**
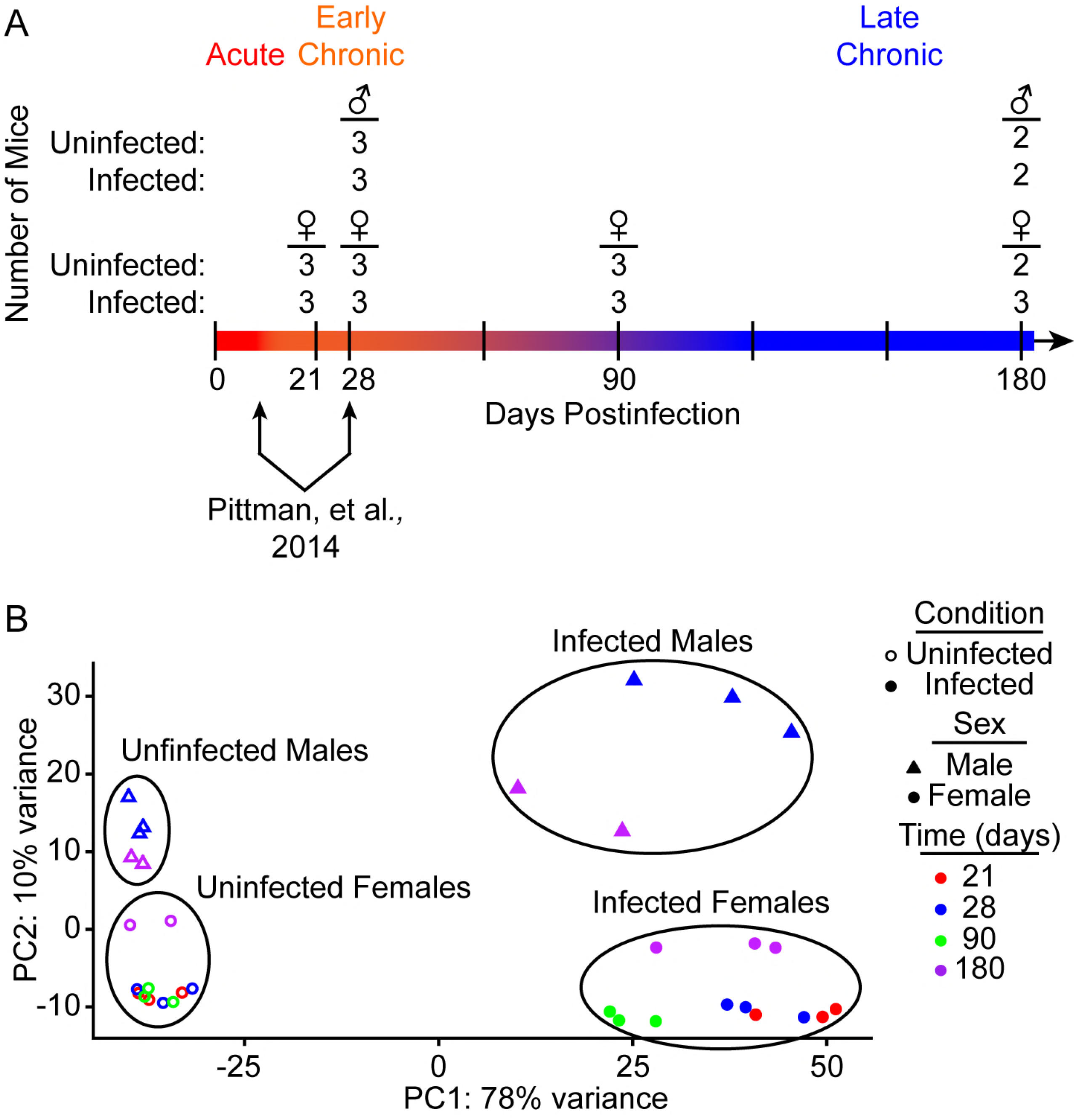
RNA-seq of long-term chronic infection. (**A**) Timeline of infection representing RNAseq timepoints, defining acute (<14 days), early chronic (21-28 days), and late chronic (>90 days) *T. gondii* infection. Female cerebral cortexes (♀) were harvested 21, 28, 90, and 180 days postinfection, and male cerebral cortexes (♂) were harvested 28 and 180 days postinfection. Numbers represent the number of individual mice used for each group in the RNAseq analysis. Arrows represent the timepoints used to compare acute and early chronic transcripts from the previous analysis (Pittman, et al., 2014). (**B**) PCA plot of samples used for RNA-seq analysis. Open symbols represent uninfected samples and closed symbols represent infected samples. Triangle represent males, and circles represent females. Colors correspond to timepoint of infection (red=21 days, blue=28 days, green=90 days, purple=180 days).

We first compared the differential expression of host genes from infected female cerebral cortex to their corresponding uninfected controls (Table S4). A statistical cutoff of an adjusted p-value (false discovery rate) of <0.05 was used for all analyses, and genes with >2-fold change from uninfected mice were identified to increase the likelihood of biological significance. This threshold identified 1660, 1369, 745, and 1256 genes differentially expressed in female mice at 21, 28, 90, and 180 days postinfection, respectively. Nearly all these genes were higher in abundance, as only 110, 28, 6, and 52 genes were less abundant at 21, 28, 90, and 180 days postinfection, respectively. This result was similar to that seen in the previous chronic transcriptional analysis of *T. gondii* infection (21). Of the significantly less abundant genes, the average change was approximately 2.5-fold for all timepoints expect for the 90-day timepoint which averaged 5.0-fold reduction. No gene was >10-fold reduced for any of the female timepoints.

### Host genes with greater abundance are shared between early and late chronic infection

Among female genes with greater abundance, 710 are shared between all timepoints (Fig. 2A), which represents approximately half of all differentially expressed genes at 21, 28, and 180 days postinfection and 95% at 90 days. The results were similar for male mice, with 609 shared genes between days 28 and 180 (Fig. 2B), out of 2009 and 1014 genes more abundant at 28 and 180 days postinfection, respectively, for males. The number of unique female genes is drastically reduced from 21 to 28 days postinfection (322 to 48, respectively). This large set of 21 day unique genes suggests these genes may be either acute stage specific genes or genes required for the onset and establishment of chronic infection. As evidence for both scenarios, 193 of the 322 genes specific for 21-days are also differentially expressed when comparing chronic transcripts at 28 days to acute transcripts at 10 days postinfection (21).

**Fig. 2.**
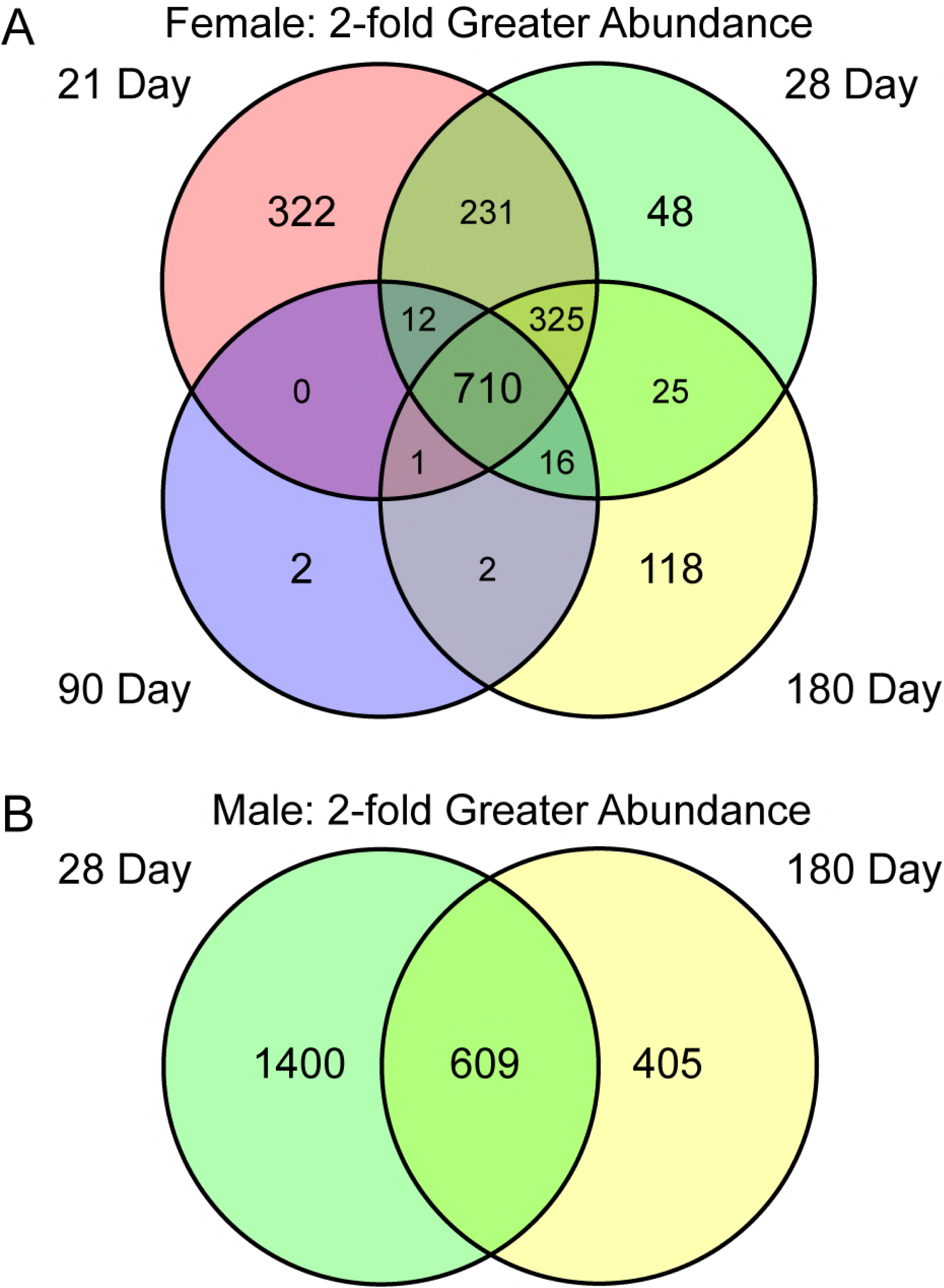
Host genes with greater abundance after infection are shared throughout infection. Number of female (**A**) and male (**B**) mouse genes with a greater than 2-fold increase from age-matched uninfected controls. Shared sets are represented by overlapping regions between the 21 day (red), 28 day (green), 90 day (blue), and 180 day (yellow) timepoints.

To understand which host genes were the most responsive to infection, we sorted for genes with a 20-fold or greater abundance in infected versus uninfected mice. 92 female genes met these criteria in both early (28-days) and late (6 months) chronic infection (Fig. 3A). Similarly, male mice have several genes (42 genes) greater in abundance both early and late into infection (Fig. 3B). 153 male genes were at least 20-fold differentially expressed during infection at 28 days, 148 of which were more abundant. At 180 days 43 male genes are differentially expressed using the 20-fold cutoff, all of which are more abundant and 42 are shared between both timepoints. Moreover, 40 of the 42 shared male genes are also shared with females at both 28 and 180 days postinfection (Fig. 3C), demonstrating that the host continues to robustly respond to *T. gondii* months after initial infection and genes with greater abundance are shared between sexes during chronic infection.

**Fig. 3.**
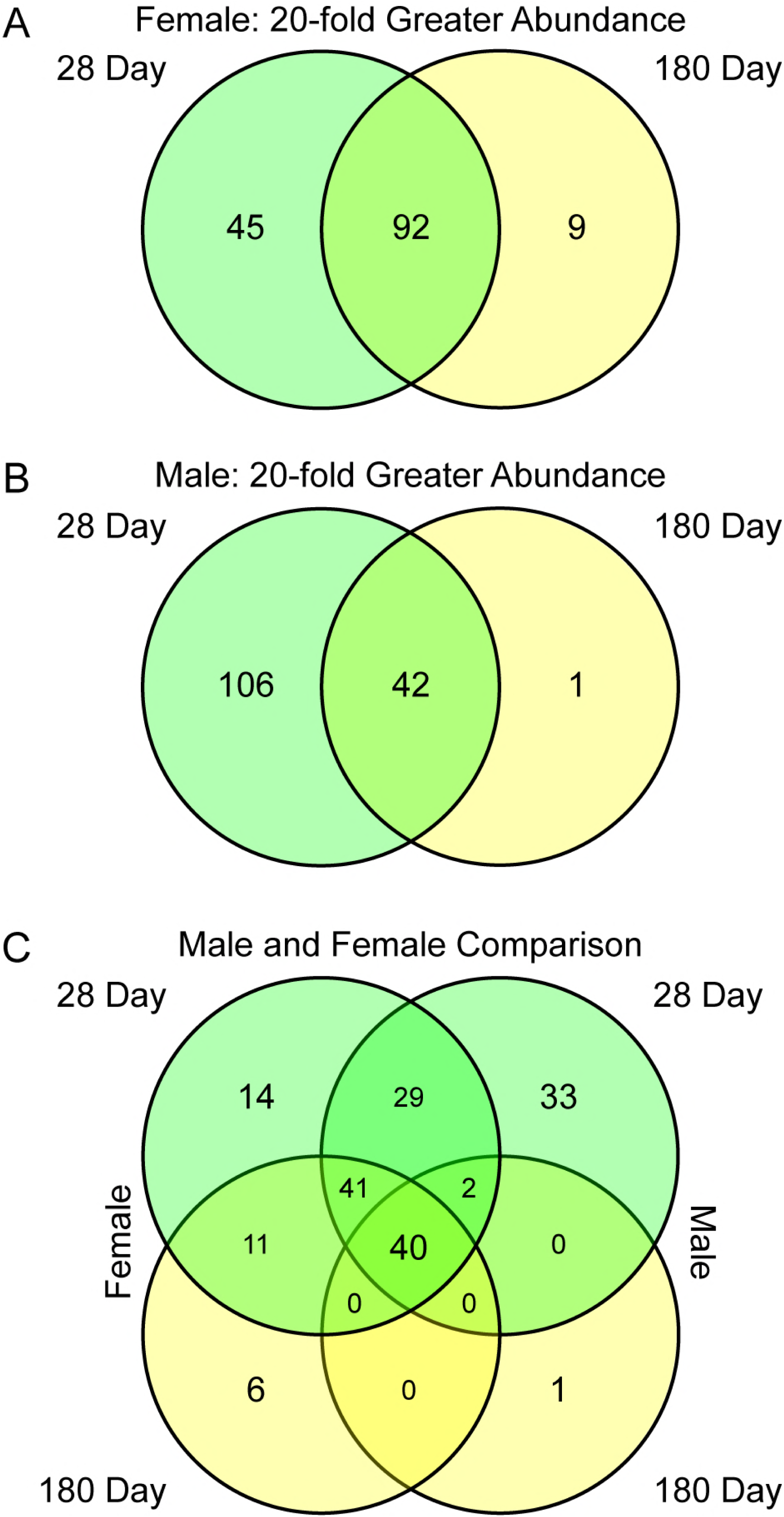
Many enriched host genes are shared throughout infection and between sexes. Number of female (A) and male (B) mouse genes with a 20-fold increase from uninfected controls. (C) Shared sets are represented by overlapping regions between the 28 day (green) and 180 day (yellow) timepoints.

### Male mice have vastly more genes that are less abundant

The number of less abundant genes was strikingly different between males and females (Fig. 4). Only 28 and 52 female genes were less abundant at 28 and 180 days postinfection, respectively. In males, 2290 genes were less abundant at 28 days and 970 at 6 months postinfection using a 2-fold cutoff, with 328 shared between both timepoints. Of these, 40 are >10-fold reduced at 28 days and 3 genes reduced at 6-months postinfection, none of which are common between the two timepoints (Table S5).

**Fig. 4.**
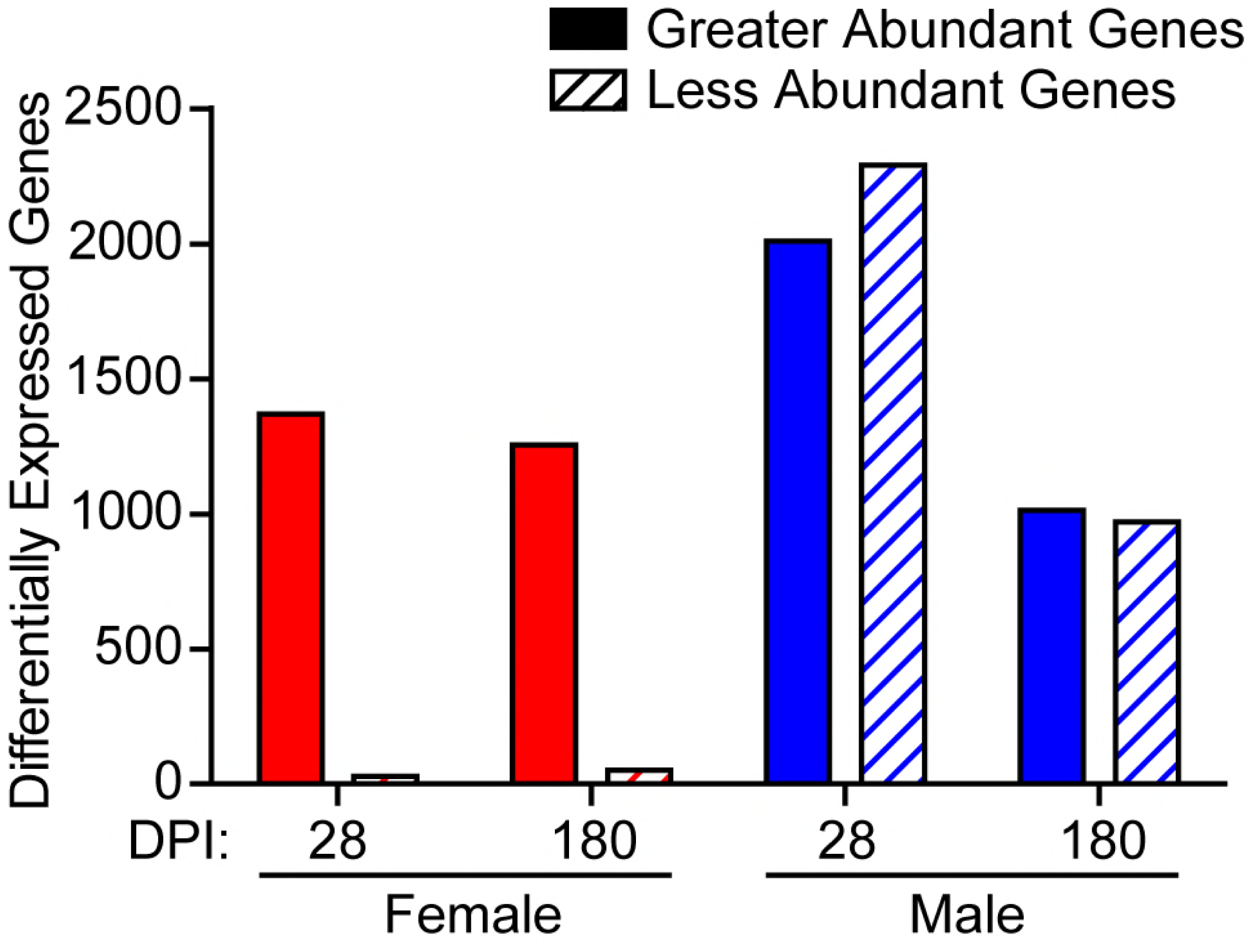
Males have more less-abundant genes than females in response to infection. Number of differentially express genes with a 2-fold increase (solid bars) or decrease (slashed bar) compared to uninfected controls for females (red) and males (blue) at 28 or 180 days postinfection (DPI).

### Gene ontology analysis shows immune response late into infection

To understand the roles for these common genes during in infection, we analyzed the sets of shared genes for gene ontology (GO) enrichment. Using the 710 greater abundant genes shared among the female timepoints, 370 GO terms were significantly enriched (Table S6). Similarly, 291 GO terms were identified within the male dataset of shared more abundant genes (Table S6). All the most enriched GO terms related to immune responses or other responses to infection and were the same for both males and females (Fig. 5A). This immunological GO term enrichment is also found among the >20-fold more abundant genes (Fig. 5B), showing that *T. gondii* continuously induces immunological responses during late chronic infection.

**Fig. 5.**
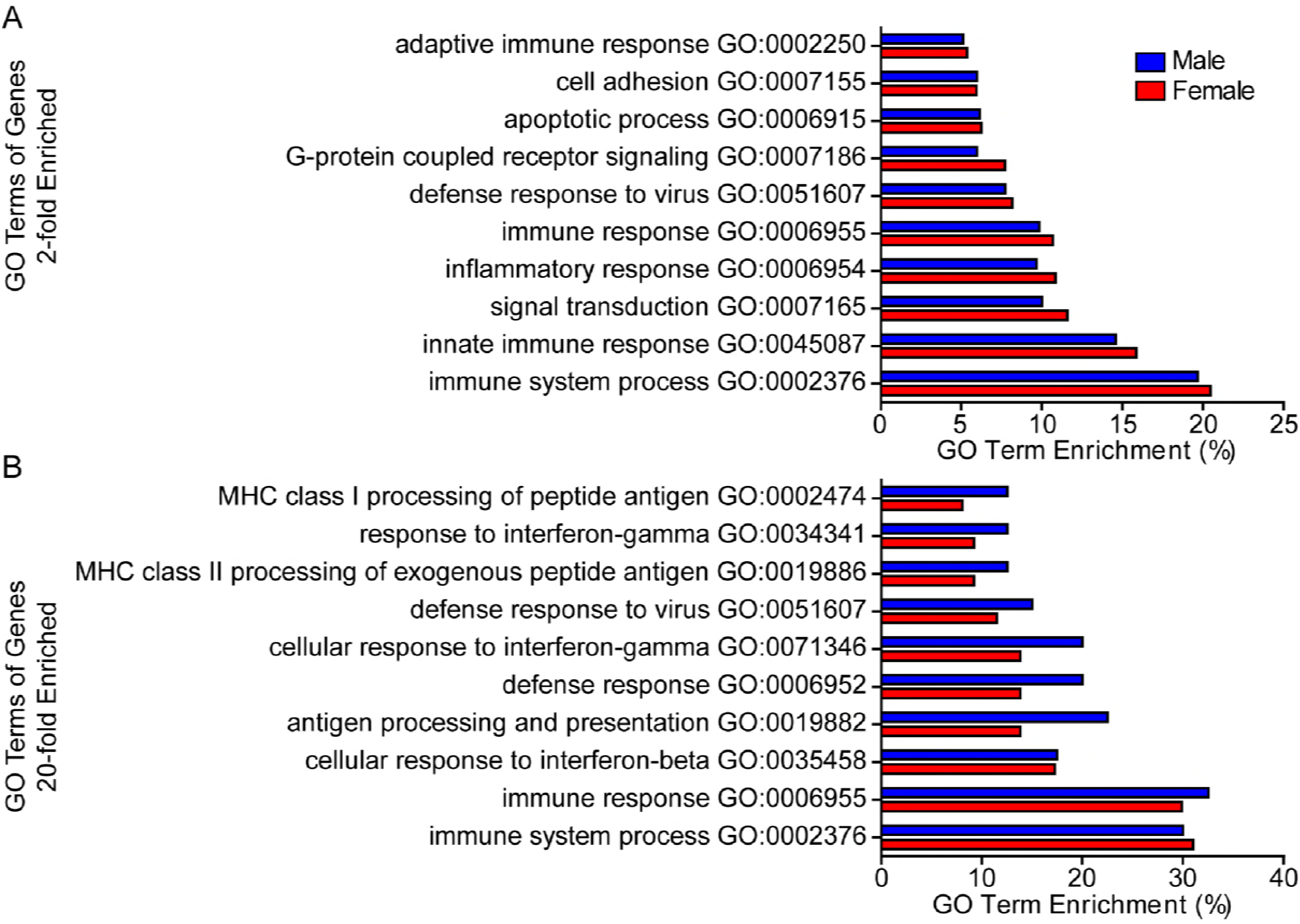
Immunological gene ontology (GO) terms are highly enriched during infection. Differentially expressed genes shared between all male (blue) and female (red) timepoints with a 2-fold (A) or 20-fold (B) increase were analyzed for GO term enrichment. Terms were sorted based on their q-value and the percent enrichment for the top ten most significant are shown.

### Serum cytokines are elevated throughout the course of infection

We quantified serum cytokines to determine whether the increased transcriptional abundance of immunological factors translated into a physiological response. Serum cytokines important for control of *T. gondii* include tumor necrosis factor alpha (TNFα), IFNγ, IL-12, IL-6, IL-10 and MCP1 (Fig. 6A-F, respectively). TNFα, IFNγ, MCP1, and IL-12 were highest in infected male and female mice at the earliest timepoint tested (21-days postinfection) and decrease as the infection continued. By 6 months postinfection, TNFα, IFNγ, MCP1, and IL-12 were all decreased by approximately 70% from the levels at 21 days postinfection; however, levels at 6-months were still significantly higher than uninfected. These results were similar to the decrease in serum cytokines seen in C57BL/6 mice at 8-weeks post *T. gondii* infection (22). While IL-6 and IL-10 both showed a trend of elevated serum concentration compared to uninfected, not all timepoints reached statistical significance. Comparing cytokines from male to female, we found the same trend of decreasing concentrations for TNFα, IFNγ, MCP1, and IL-12 for both sexes. In addition, IL-6 and IL-10 were variably significant for both sexes throughout infection compared to uninfected. Although some cytokines tended to be lower in males, these differences were not statistically different.

**Fig. 6.**
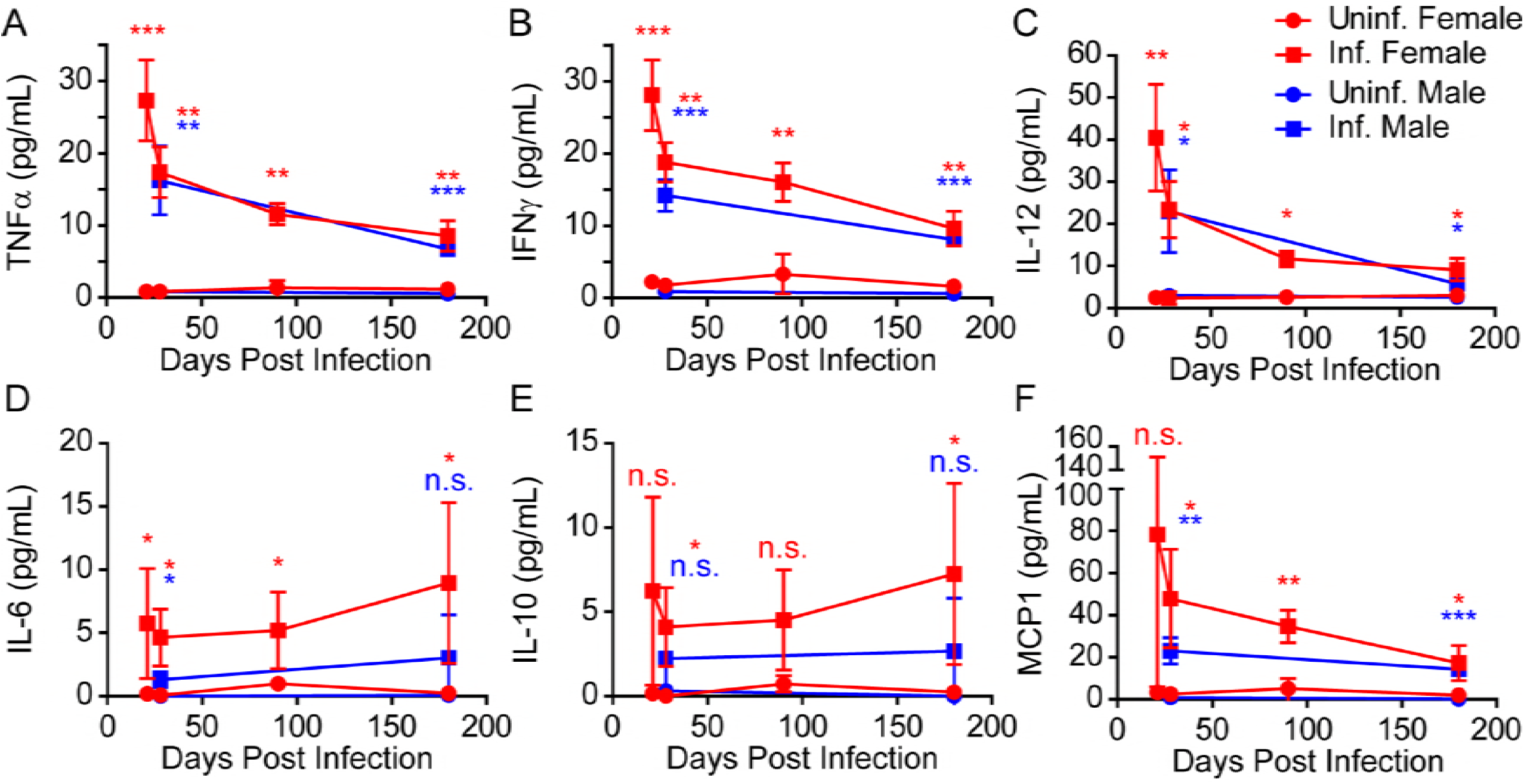
Inflammatory cytokines are abundant throughout infection. Serum TNFα (A), IFNγ (B), IL-12 (C), IL-6 (D), IL-10 (E) and MCP1 (F) were quantified from the male (blue) and female (red) mice used for the RNAseq analysis. Timepoints include 21, 28, 90 and 180 days postinfection. Circle symbols represent cytokines from infected mice and squares symbols represent uninfected mice. Data is the average concentration of three mice, except for infected and uninfected males at 180 days postinfection and uninfected females at 180 days postinfection, which had two mice each. Error bars represent the standard deviation between samples and asterisks indicate statistically significant differences between infected and uninfected samples by Student’s t-test (n.s., not significant; *, p<0.05; **, p<0.01; ***, p<0.001).

The cytokine concentrations matched the transcriptional profile over time for many of the tested cytokines for both sexes. The normalized expression values (FPKM values) for TNFα, IFNγ, and IL-12 had the highest levels in the infected mice at 21 days postinfection and decreased over time. Many immunological genes show a dip in transcriptional activity at 90 days postinfection, but the transcriptional change did not correlate to a reduction in serum cytokines tested. Serum concentrations of MCP1 were elevated even though mRNA levels of MCP1 in the cerebral cortex were not significantly changed from uninfected at any of the timepoints. This result suggests that MCP1 is not produced by glial cells in response to chronic *T. gondii* infection, but instead this response is initiated outside of the brain. Gene expression of IL-6 and IL-10 showed low and unchanging levels throughout infection, and these levels were not significantly different from uninfected mice at any of the timepoints in the female mice. Expression for the receptors for many of the tested cytokines are increased in abundance, suggesting the host brain cells are responsive and active toward the immunological cytokine response.

### Sex specific responses to infection

Because male and female mice had similar immunological GO term enrichment in response to *T. gondii* infection, we expected their survival to *T. gondii* would also be similar. During an acute challenge with 1×10^5^ parasites/mouse, both male and female mice became moribund and required euthanasia at similar rates (Fig. 7A). When mice are infected with a lower dose of parasites (1×10^4^ parasites/mouse) to allow survival into chronic infection, both sexes had 90% survival at the start of chronic infection (28 days) (Fig. 7B). Although male mice had a slightly lower survival percentage throughout chronic infection, the survival curve did not reach statistical significance between sexes. Males had a 2-fold higher cyst at 3 months postinfection (Fig. 7C). As the majority of the mice do not survive to 6 months postinfection, the cyst counts in the remaining mice are highly variable, especially in the males (Fig. 7C). These small differences in survival and cyst counts may be related to the slight reduction in key cytokines in males during chronic infection (Fig. 6).

**Fig. 7.**
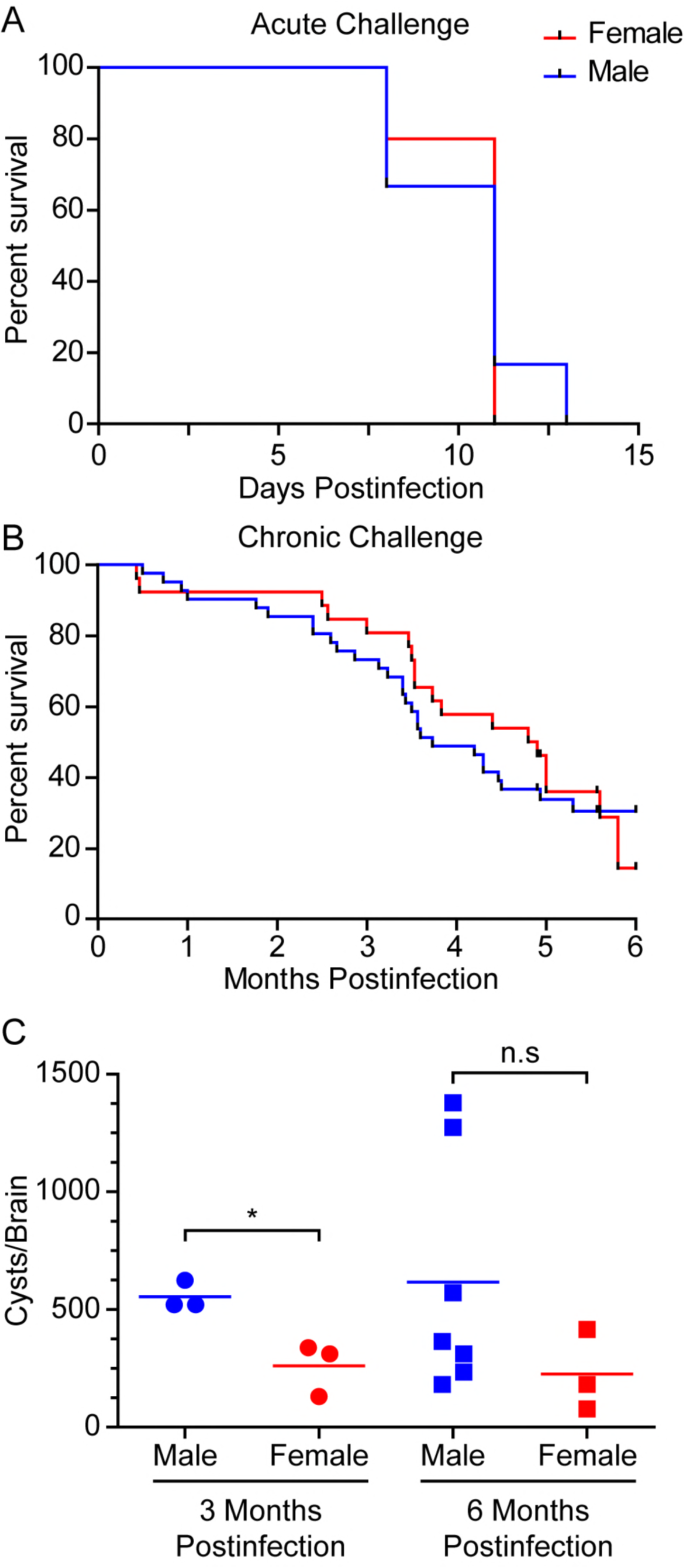
Outcomes of *T. gondii* infection are similar in male and female mice. (A) Acute survival curve of male (blue, n=6) and female (red, n=5) mice infected with 1×10^5^ parasites. (B) Chronic survival curve for male (n=28) and female (n=26) mice infected with 1×10^4^ parasites. (C) Cyst burden in brains of mice at 3 months (n=3 male, n=3 female) and of surviving mice at 6 months postinfection (from **B**; n=7 for males and n=3 female). Statistical differences between male and female survival curves was calculated using the Mantel Cox log-rank test, and differences between male and female cyst burdens was determined using Student’s t-test (n.s., not significant; *, p<0.05).

Although most differentially expressed genes were common between sexes, 11 genes (Table S7) were increased 20-fold in females but not in males (Fig. 3C). The two genes with the greatest difference between sexes were CXCL13 and CCR2. CCR2 is the receptor for MCP1, and *T. gondii* PRF confers resistance to a secondary bacterial infection by recruiting monocytes in a CCR2 dependent manner (8). We tested the ability of *T. gondii* to confer resistance to *Listeria monocytogenes* in male mice during chronic infection (Fig. 8A) and found a 3.3 log reduction in CFU from the spleens of male mice chronically infected with *T. gondii*. This was similar, to the 3.6 log reduction seen in chronically infected females (8). To directly compare male and females, and as PRF is the CCR2 stimulant, we treated mice with 10 ng PRF (as a proxy for chronically infected mice) prior to *L. monocytogenes* infection (Fig. 8B). The 3.2 log CFU reduction seen in male mice when treated with PRF was nearly identical as chronic infection, and female mice showed a 3.6 log reduction, similar to results seen previously (8). Although female mice had an overall higher CFU burden compared to males, the CFU reduction was slightly greater for female mice, but this difference was not statistically different. While female and male mice have similar protection against *T. gondii* (Fig. 7) and *L. monocytogenes* (Fig. 8), the precise molecules and their intensities of that protection are different, so mechanistic analyses should include both sexes in the studies.

**Fig. 8.**
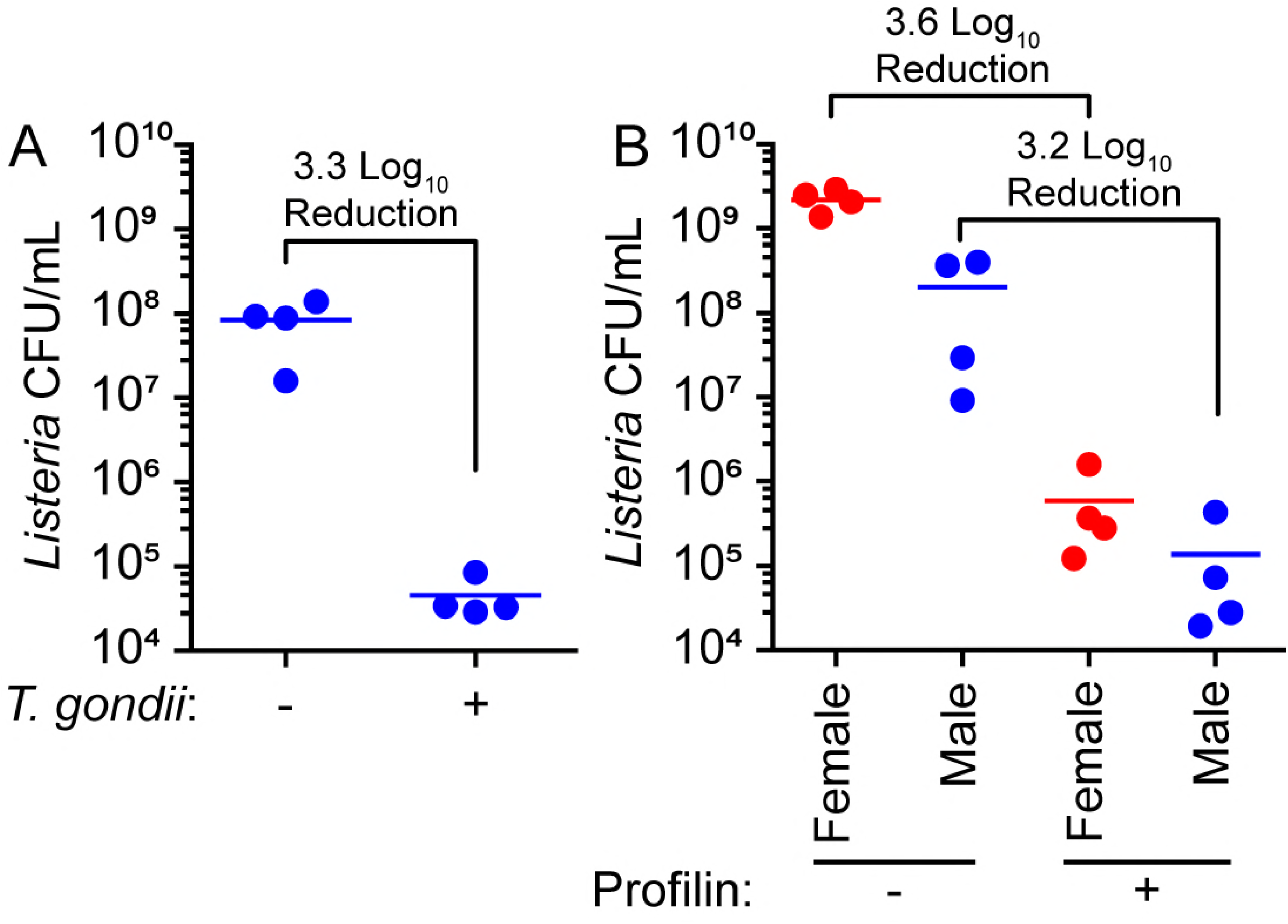
Stimulation with *T. gondii* confers more resistance to *L. monocytogenes* in female mice. (**A**) *L. monocytogenes* burden in spleens of male mice infected 28 days prior to *L. monocytogenes* with *T. gondii* (+, n=4) compared to uninfected male mice (-, n=4). (**B**) *L. monocytogenes* burden in spleens of female (red) and male (blue) mice (n=4/group) treated intravenously with saline (-) or 10 ng of recombinant *T. gondii* PRF (+) 4 hours prior to *L. monocytogenes* infection. Lines represent mean burden of each group.

## DISCUSSION

*T. gondii* can form a chronic infection in any warm-blooded animal, presumably for the life of the host. In mice, this chronic infection alters the host both physically and behaviourally. Mice show increased activity relative to uninfected controls (23, 24) and infected mice are no longer repulsed by feline urine (25). This type of host manipulation serves as a selective advantage for the feline/rodent cycle. However, the induced behavioural changes and their effects in humans is more controversial and less understood. Associations with *T. gondii* in humans range from personality changes, to increased risk of involvement in a traffic accident, to mental health disorders such as schizophrenia (26, 27). Because no drug treatments are effective against chronic *T. gondii* infection and there are no effective vaccine options, it is important for us to better understand the long-term effects on the host. Our current RNA-seq study of a long-term chronic infection time course will enhance our understanding of these behavioural studies and their effects on the brain.

Our transcriptional analysis of mouse genes during late chronic infection shows a continuous and robust immunological response to *T. gondii*. Host responses are reduced after the 21 day timepoint, but they are still significantly higher than the age-matched uninfected mice. Nearly half of all the genes that increased in abundance at 21 days post infection (start of chronic infection) were still increased 6 months postinfection (Fig 2). All the most enriched GO terms from this set of genes related to immunological responses (Fig. 5). Although the amplitude of expression for immunological genes during late-chronic infection may be reduced for some genes such as TNFα and IFNγ, they are still significantly more abundant than in the uninfected controls and many genes are >20-fold increased (Fig. 3). Given these results, it is not surprising that natural killer cells isolated from the peritoneal cavity of mice continue to be activated out to 6 months postinfection (28) and *T. gondii* infection maintains protection from secondary infection out to 7-months postinfection (2). This demonstrates a continual immunological response from the host and has been verified in our transcriptional analysis here.

Both sexes showed a major increase in the expression of several immunological genes. While the majority of the differentially expressed genes are shared, key factors showed differences between the sexes leading to altered responses to the infection. Levels of CCR2 expression was higher in females, along with a trend of increased serum cytokine concentrations, compared to male mice (Fig. 6). We previously found that CCR2 is essential for *T. gondii* chronic infection and PRF to confer resistance to *L. monocytogenes* in females but these findings were not tested male mice. In this current study, we found that both chronic infection and PRF treatment were effective to reduce the *L. monocytogenes* bacterial burden in male mice, similar to female mice. PRF treatment reduced bacterial burden in males by 3.2 logs compared to untreated males, related to the bacterial reduction of 3.6 logs for females (Fig. 8). These results emphasize that while there are some differences in the molecules induced by *T. gondii*, *T. gondii* can protect against *Listeria* in both sexes. Future immune mechanistic analyses should include both female and male mice.

A striking difference between the sexes was seen in the number of genes with lowered abundance during infection. While female mice only had around one hundred genes with reduced abundance at each timepoint, male mice had over 2000 genes at 28 days postinfection and over 1000 at 6 months postinfection with lowered abundance (Fig. 4). Many of the most reduced genes have an unknown function (Table S4), but five of the most reduced genes are named genes: FCRLS, DLG2, PPIL6, SCEL, and MARCH10. These genes are all reduced specifically in males except for FCRLS which is reduced at in both sexes. Disks large homolog 2 (DLG2, transcript number ENSMUST00000207095) is a membrane-associated guanylate kinase that forms multimeric scaffolds at postsynaptic sites. While DLG2 knockout mice have normal synapse development (29), they have abnormal thermogenesis and bone density (30), suggesting male mice may have altered metabolism during *T. gondii* infection. Peptidylprolyl isomerase like 6 (PPIL6), is a gene enriched in olfactory sensory neurons (31); however, the biological function is unknown, as the PPIL6 catalytic site is inactive (32). As mice have altered perception of feline urine scent (25), PPIL6 may play a role in this mechanism for male mice.

Sciellin (SCEL), is a protein thought to function in the assembly or regulation of cornified envelope proteins in keratinocytes (33), and membrane associated ring-CH-type finger 10 (MARCH10) is a ubiquitin ligase (34). Although the role of SCEL and MARCH10 in the brain has not been determined, links of these genes can be drawn to Alzheimer’s disease. A specific type of keratin has been detected in cerebrospinal fluid of patients with Alzheimer’s disease but not healthy individuals, which may be the result of blood brain barrier breakdown or dysregulation of the ubiquitin proteasome (35). Since SCEL is involved with cells which produce keratin and MARCH10 is a ubiquitin ligase, these genes may play a role in the mechanism of keratin accumulation in the CNS.

One host gene less abundant in both males (28-fold at 28 days) and females (8-fold at 28 days) is Fc receptor-like S scavenger receptor (FCRLS), a protein with unknown function that is highly expressed on microglia (36). FCRLS resembles an Fc receptor but does not bind antibodies. It is interesting to notice that other microglial-specific markers, such as P2ry12, Mertk and Gas6, have similar transcript levels in infected and uninfected males, while C1qa and Pros1 are significantly more abundant in infected versus uninfected males (Table S3). These results point to a specific downregulation of FCRLS in microglia by *T. gondii*, and not a global downregulation or lysis of microglia cells by *T. gondii*.

This current study is another example of the importance of using male and female mice when studying the immune responses to *T. gondii*. Our previous studies of chronic *T. gondii* infection being protective against other pathogens were all performed exclusively in female mice (8, 37, 38). Our methods in the manuscript did not state the sex of the mice, but our records indicated that only female mice were used. The original studies on the effects of chronic infection being protective against other pathogens either did not say the sex of the animal (4) or they only used female mice (5–9), so this study is the first to show that the protective effects of *T. gondii* chronic infection are not exclusive to female mice. This phenomenon and several other immune studies will have to be re-examined in both male and female mice in light of these new findings to better understand the host immune response to *T. gondii*.

## METHODS

### Mouse infection with T. gondii

The ME49 strain of *T. gondii* was cultured as tachyzoites in human foreskin fibroblast cells. 6-10 week-old CBA/J male and female mice (JAX) were infected intraperitoneally with 1×10^5^ tachyzoites for studies of acute toxoplasmosis and 1×10^4^ tachyzoites to establish chronic toxoplasmosis. Animal survival curves were produced and analyzed using Prism software (v5; GraphPad). Animals were treated in compliance with the guidelines set by the Institutional Animal Care and Use Committee (IACUC) of the University of Wisconsin School of Medicine and Public Health (protocol #M005217), which adheres to the regulations and guidelines set by the National Research Council. The University of Wisconsin is accredited by the International Association for Assessment and Accreditation of Laboratory Animal Care.

### Generation of RNA and RNAseq

The cerebral cortex of infected mice at 21, 28, 90, and 180 days postinfection, along with aged-matched uninfected control mice, was harvested. Brain tissue from three mice were processed for RNAseq for each condition and sex for the 21, 28, and 90 day timepoints, and two mice were used for each condition for the 180 day timepoint, except the infected female 180 day timepoint group in which three mice were used. Tissue was immediately homogenized in Trizol (Ambion) using a pellet pestle (Kontes), and RNA extracted using phenol/chloroform separation and ethanol precipitation. RNA libraries were prepared using TruSeq RNA Library Prep Kit v2 set A (Illumina), according to the manufacturers protocol. RNA libraries were multiplexed and run across 9 lanes (infected tissues) or 3 lanes (uninfected tissues) for paired end sequencing by Illumina HiSeq 2500 at the University of Wisconsin Biotechnology Center, generating approximately 5.7 billion total paired-end 125 base pair reads.

### Differential Expression Analysis

The sequenced reads were aligned to the *Mus musculus* genome (GRCm38; database v87; Ensembl), using the Spliced Alignment to a Reference (STAR; v2.5.2b) program (39). Alignment parameters for STAR were kept at default except for the following: 2 base pair maximum mismatch, 70 base pair minimum intron length, and a 500,000 base pair maximum intron length. Gene expression levels were then estimated using the program RNA-seq by Expectation-Maximization (RSEM v1.2.31) (40). Differential analysis was calculated using the DESeq2 (v1.14.1) program, analyzing genes with ≥10 raw counts (calculated by RSEM) for at least one sample. Fold change was calculated using the geometric mean values for the infected samples relative to their matching uninfected samples. The rlog transformed DESeq2 values were used for the PCA plot and PCA calculated using “plotPCA” function of the DESeq2 package. The log_2_ transformed DESeq2 values were used to calculate Pearson’s correlation coefficients and coefficients were calculated using the corrgram package. The Database for Annotation, Visualization and Integrated Discovery (DAVID v6.8) was used as a functional annotation tool to identify enriched biological gene ontology (GO) terms among genes commonly differentially expressed between timepoints. A list of mouse genes with at least 10 RSEM gene counts was used as background for enrichment analysis in DAVID. All raw sequencing data and differential expression values have been deposited in NCBI’s Gene Expression Omnibus (GEO) (41) and are accessible through GEO Series accession number GSE117504 (https://www.ncbi.nlm.nih.gov/geo/query/acc.cgi?acc=GSE117504).

### Listeria monocytogenes infection

C57BL/6 mice were either infected peritoneally with 250 ME49 tachyzoites or treated retro-orbitally with 10 ng *T. gondii* PRF (8, 10) or PBS as control treatment. Purified recombinant his-tagged *T. gondii* PRF was a kind gift from F. Yarovinsky which consistently yields near endotoxin-free protein (10). Mice were infected with approximately 6×10^4^ *L. monocytogenes* colony forming units (CFU) retro-orbitally either 28 days post *T. gondii* infection or four hours post-PRF treatment. Bacterial culture was grown to mid-log growth stage at 37°C in LB medium prior to infection. At 3 days post *L. monocytogenes* infection, spleen tissue was harvested and homogenized in 1 mL PBS using a pellet pestle (Kontes). Homogenate was plated on LB agar plates to quantify viable *L. monocytogenes* CFU for each condition.

### Quantitation of brain cyst burden

Whole brain tissue was homogenized using a pellet pestle (Kontes) in PBS. Homogenate was fixed in 4% paraformaldehyde, then blocked and permeabilized in PBS with 3% BSA and 0.2% Triton X-100. Biotinylated *Dolichos biflorus* agglutinin (DBA; Vector Laboratories) was used to mark the surface of the *T. gondii* cyst and was visualized using Alexaflour-594 conjugated streptavidin (Thermo Scientific). Cysts were counted by microscopy to calculate cyst burden for each brain.

### Quantitation of serum cytokines

Blood was harvested from the mice used for RNAseq analysis via cardia puncture. Samples were incubated on ice, and serum was separated from the coagulated clot by centrifugation (10 minutes at 14000xg). 25 µL of serum was used for cytokine quantification using a cytometric bead array (BD Biosciences). Serum cytokine concentrations were calculated based on a standard curve of known concentrations.

## ACKNOWLEDGEMENTS

We would like to thank Charles Czuprynski for *L. monocytogenes* EGD strain and Felix Yarovinsky for recombinant *T. gondii* profilin. This research was supported by the National Science Foundation fellowship DGE-1747503 (P.W.C), National Institutes of Health (NIH) National Research Service Award T32AI007414 (P.W.C) and T32AI55397 (A.L.G), Science and Medicine Graduate Research Scholars program (P.W.C), and NIH R21AI114277 (L.J.K.).

